# Diagnostic value of galactomannan in bronchoalveolar lavage fluid for chronic respiratory disease with pulmonary aspergillosis

**DOI:** 10.1101/730770

**Authors:** Guangbin Lai, Chao Zeng, Jianming Mo, Wei-dong Song, Ping Xu

## Abstract

**Objective:** To explore the diagnostic value of the bronchoalveolar lavage fluid galactomannan (BALF GM) test for chronic respiratory disease with pulmonary aspergillosis, and establish the optimal cut-off value.

**Methods:** A total of 180 chronic respiratory disease patients seen at the Respiratory Medicine Department of Peking University Shenzhen Hospital from September 2017 to September 2018 were analyzed. According to the diagnostic criteria, we divided the patients into the case group (n = 70, comprising 5, 20, and 45 proven, probable, and possible cases, respectively) and the control group (n = 110). Bronchoalveolar lavage fluid was collected, and the BALF GM test results were analyzed.

**Results:** A non-parametric rank-sum test showed that the mean rank of the case group was 140.80, which was higher than that of the control group (58.49). The Z-value was 10.335 (P = 0.000), indicating that the general distribution of BALF GM differed between the two groups. A BALF GM cut-off value of 0.485 showed the highest diagnostic efficacy for pulmonary aspergillosis. The sensitivity, specificity, positive predictive value, and negative predictive value were 92.9%, 100%, 92.8%, and 100%, respectively. As the cut-off value increased, the specificity and sensitivity of the BALF GM test increased and decreased, respectively.

**Conclusions:** The BALF GM test can be used confirm the diagnosis of patients with pulmonary aspergillosis and chronic respiratory disease. The optimum BALF GM cut-off value was 0.485. Antifungal therapy is important for treating pulmonary aspergillus infection in patients with chronic respiratory disease.

---

Pulmonary aspergillosis (PA) is common in patients with severe immunosuppression, especially recipients of hematopoietic cell transplantation (HCT) or solid organ transplantation (particularly the lung, cardiopulmonary system, and liver). As well as patients with a long history of neutropenia, individuals with some hereditary immunodeficiencies (such as chronic granulomatous disease) also have a higher risk of pulmonary aspergillosis^[1]^. Invasive pulmonary aspergillosis (IPA) can also occur in individuals with less severe or no immunosuppression, especially in those with potentially chronic respiratory diseases^[2, 3]^. The situation is particularly severe in South China with a humid and warm subtropical climate. According to the World Health Organization (WHO) definition^[4]^, chronic respiratory disease refers to a group of chronic diseases that affect the airways and other structures of the lungs. The most important of these are chronic obstructive pulmonary disease (COPD), bronchial asthma, and bronchiectasis.

Chronic respiratory disease with pulmonary aspergillosis is occult and has a high mortality rate. In the retrospective study by Guinea *et al*.,^[5]^ *Aspergillus* was isolated from 239 of 14,618 patients with COPD, pulmonary aspergillosis was diagnosed in 53 patients, 38 of whom died within 4 months of diagnosis (mortality rate, 71.6%). Early diagnosis of this type of disease is particularly important, and the bronchoalveolar lavage fluid galactomannan (BALF GM) test is a useful auxiliary diagnostic modality. Indeed, in 2016, Infectious Diseases of America (IDSA) recommended BALF GM as a diagnostic marker for aspergillosis^[6]^. However, there is no consensus on GM reference value in alveolar lavage fluid, not only in immunodeficiency patients, but also in patients with chronic airway diseases. In order to evaluate whether the GM of alveolar lavage fluid be a diagnostic marker for patients with chronic airway diseases complicated with pulmonary aspergillosis, and to find out the appropriate GM cut-off value, We studied patients with COPD, bronchial asthma, bronchiectasis and other chronic respiratory diseases commonly seen in the Department of Respiratory Medicine, Peking University Shenzhen Hospital. We collected BALF and evaluated the utility of the BALF GM level as a diagnostic marker of, and treatment target for, chronic respiratory diseases with pulmonary aspergillosis.

## 1 Materials and Methods

### 1.1 Enrollment

A total of 180 patients admitted to hospital from September 2017 to September 2018 due to chronic respiratory diseases (COPD, bronchial asthma, or bronchiectasis) were enrolled based on the following inclusion criteria: (1) risk of pulmonary infection (imaging examination revealed the presence of lung lesions); (2) no contraindications for bronchoscopy and provision of informed consent; and (3) no antifungal use before admission. The study protocol was approved by the Ethics Committee of our hospital.

### 1.2 Diagnostic classification

According to the 2016 IDSA guidelines on *Aspergillus* infection diagnosis and treatment ^[6]^, together with the diagnostic criteria for invasive fungal infections used in China^[7]^, the cases of chronic respiratory disease with pulmonary aspergillosis were classified as follows:

(1) Proven (confirmed diagnosis): a lower respiratory tract specimen positive for *Aspergillus*, with confirmation by molecular, immunological, and/or culture methods that the observed hyphae were of *Aspergillus*.

(2) Probable (clinical diagnosis): frequent use of hormones or repeated use of broad-spectrum antibiotics for 3 months; suggestive pulmonary imaging changes within 3 months; and positive for *Aspergillus* or a positive result of a blood or BALF GM test.

(3) Possible: patients with chronic respiratory diseases who frequently use hormones or have been treated with broad-spectrum antibiotics for 3 months, and who recently developed acute exacerbations and presented with indicative changes in lung images within 3 months (chest X-ray or computed tomography [CT]), but in whom *Aspergillus* was not cultured from lower respiratory tract specimens and the results of microscopic and serological tests were negative.

(4) *Aspergillus* colonization: lower respiratory tract specimens of patients with chronic respiratory diseases positive for *Aspergillus*, but without acute exacerbation, airway spasm, or new pulmonary infiltration.

### 1.3 Group assignment

total of 180 patients with chronic respiratory diseases were enrolled (92 males and 88 females), comprising 58 patients with chronic obstructive pulmonary disease, 59 with bronchial asthma, and 63 with bronchiectasis. According to the clinical symptoms and the results of auxiliary examinations, the patients with proven, probable, or possible pulmonary aspergillosis were included in the case group according to the above-described diagnostic standards (n = 70; 5, 20, and 45 proven, probable, and possible cases, respectively). Patients with non-pulmonary aspergillosis constituted the control group (n = 110). There was no significant difference in basic characteristics between the case and control groups (all, P > 0.05; Table 1).

**Table 1.** Basic characteristics of the case and control groups.

| Project | Case group | Control group | Statistics* | p value |
| --- | --- | --- | --- | --- |
| Number of cases (n) | 70 | 110 | - | - |
| Sex: Male,%) | 33 (47.1) | 59 (53.6) | 0.722 | 0.396 |
| Basic information (x ± s) |  |  |  |  |
| Age | 53.89±14.57 | 52.26±14.39 | -0.734 | 0.464 |
| Hospital stay (d) | 8.79±4.24 | 8.51±3.19 | -0.498 | 0.619 |
| White blood cell count (×10<sup>12</sup>/L) | 7.29±3.09 | 6.88±2.51 | -0.963 | 0.337 |
| Number of platelets (×10<sup>9</sup>/L) | 256.6±81.05 | 261.57±80.92 | 0.401 | 0.689 |
| Hemoglobin (g/L) | 129.1±17.80 | 130.71±16.72 | 0.614 | 0.540 |
| Total protein (g/L) | 67.56±6.77 | 67.72±7.51 | 0.151 | 0.880 |
| Albumin (g/L) | 37.63±5.24 | 38.51±5.01 | 1.132 | 0.259 |
| Cooccurring chronic respiratory diseases (%) |  |  |  |  |
| COPD | 23 (32.85) | 35 (31.81) | 0.021 | 0.884 |
| Bronchial asthma | 22 (31.43) | 37 (33.64) | 0.095 | 0.758 |
| Bronchiectasis | 25 (35.72) | 38 (34.55) | 0.026 | 0.873 |

\* Count data were analyzed by X<sup>2</sup>; all other data were analyzed by t-test.

### 1.4 Collection and processing of BALF

#### 1.4.1 Collection of BALF

According to the results of chest CT, bronchoscopy and bronchoalveolar lavage were performed without antifungal therapy. Bronchoalveolar lavage examinations were performed according to the expert consensus,^[8]^ as follows. (1) Site selection: according to lesion severity, the right middle or left upper lobe was selected in patients with diffuse lesions in both lungs^[9, 10]^. (2) Injection of saline:First, the bronchoscope was embedded in the opening of the bronchial segment or subsection at the lavage site. Next, at the bronchial segment or sub-segment opening, 60–80 mL of physiological saline was injected as a 20 mL bolus, at 37°C or room temperature. (3) Vacuum suction: 10–15 s after infusion of normal saline, BALF was obtained at a negative pressure of < 100 mmHg (total recovery, > 50%). Five milliliters of BALF were used for GM testing and a further 5 mL for fungal culture and other tests. (4) Finally, bronchial biopsy, lung biopsy, or bronchial brushing was performed according to the characteristics of the lesion.

#### 1.4.2 BALF GM assay

GM in BALF was quantified by sandwich assay using an antibody against *Aspergillus* GM. The absorbance value was determined in accordance with the instructions of the *Enzyme-linked Immunosorbent Assay Kit for Aspergillus*, and the galactomannan index (GMI) value was calculated. Negative and positive controls were performed in each experiment.

### 1.5 Data analysis

Data were analyzed using SPSS software (ver. 23.0; SPSS Inc., Chicago, IL, USA). Measurement data are presented as means ± standard deviation, and count data as percentages or numbers of cases. The data were tested for normality; normally distributed data were evaluated by independent sample *t*-test and chi-squared test, and non-normally distributed data were subjected to non-parametric tests. The diagnostic efficacy of BALF GM was evaluated by calculating the sensitivity, specificity, positive predictive value (PPV) and negative predictive value (NPV). Finally, a receiver operating characteristic (ROC) curve was constructed to determine the optimum cut-off value of BALF GM.

## 2 Results

### 2.1 GM status and pathogens in BALF in the case group

Using a GM cut-off value of 0.5, the GM status and pathogens detected in the case group are shown in Table 2. The BALF GM test had a higher positive rate than the serum GM test, and BALF pathogen culture showed some diagnostic utility.

**Table 2.** Cases positive for BALF GM, serum GM, and pathogens.

| Grouping | Number of positive cases (%) | Number of negative cases (%) | Total cases (%) |
| --- | --- | --- | --- |
| BALF GM test | 65 (92.8%) | 5 (7.2%) | 70 |
| Serum GM test | 7 (10%) | 63 (90%) | 70 |
| BALF pathogen culture | 15 (21.4%) | 55 (78.6%) | 70 |

### 2.2 BALF GM

By non-parametric rank-sum test, the mean rank of the case group was 140.80, which was higher than that of the control group (58.49) (Table 3). The BALF GM value of the case group was higher than that of the control group. Mann-Whitney U and Wilcoxon W tests (Table 4) yielded a Z-value of -10.335 (P = 0.000), the general distribution of BALF-GM in the two groups was different.

**Table 3.** Results of BALF GM tests.

| Group | Number of cases | Average rank | Rank sum |
| --- | --- | --- | --- |
| Control group | 110 | 58.49 | 6,434.00 |
| Case group | 70 | 140.80 | 9,856.00 |
| Whole cohort | 180 |  |  |

**Table 4.** Results of the statistical analysis.

|  | BLAF-GM test |
| --- | --- |
| Mann-Whitney U test | 329.000 |
| Wilcoxon W test | 6,434.000 |
| z value | -10.335 |
| p-value | 0.000 |

### 2.3 Diagnostic efficacy of the BALF GM test

We constructed a ROC (Figure 1) and evaluated the diagnostic efficacy of BALF GM for chronic respiratory disease with pulmonary aspergillosis, by calculating the Youden index and the area under the curve (AUC; Tables 4 and 5). The diagnostic efficacy was highest with a BALF GM cut-off value of 0.485, for which the sensitivity, specificity, PPV, and NPV were 92.9%, 100%, 92.8%, and 100%, respectively. Sensitivity decreased, and specificity increased, as the BALF GM cut-off value increased.

**Table 5.** AUC values for the BALF GM test.

| Test | Area under the curve | Standard error | Significance | Asymptotic 95% confidence interval |  |
| --- | --- | --- | --- | --- | --- |
|  |  |  |  | Lower limit | Upper limit |
| BALF GM | 0.957 | 0.021 | 0.000 | 0.916 | 0.999 |

**Figure 1.**
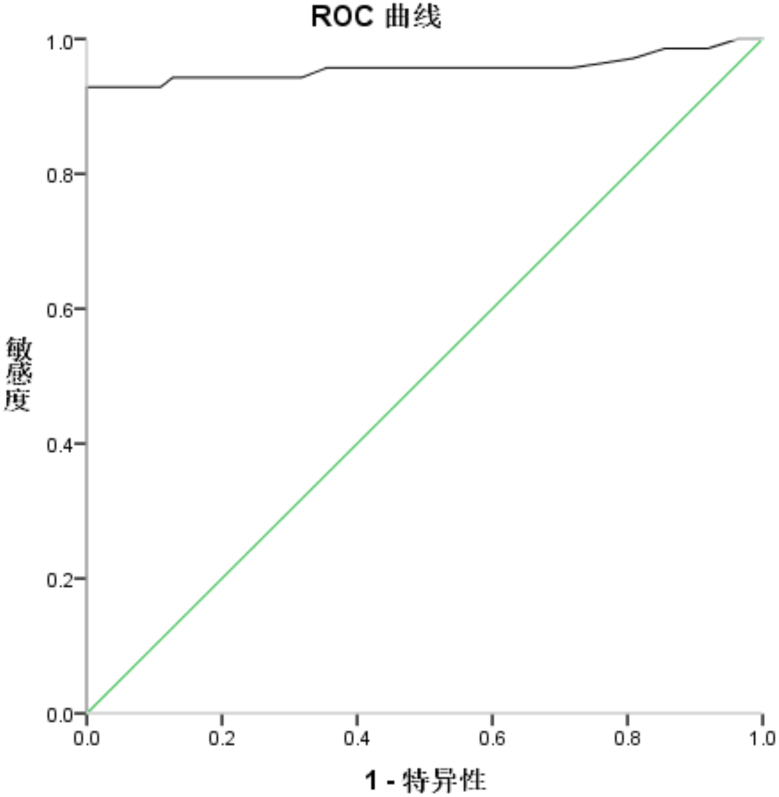
Receiver operating characteristic (ROC) curve.

### 2.4 Treatment status of patients in the case group

Seventy patients in the case group received antifungals, such as voriconazole and fluconazole. After treatment with antifungals, 20 cases were cured and 40 were improved (efficacy rate, 86%). Ten cases experienced adverse effects (adverse event rate, 14%), one of whom died (Table 7).

**Table 6.**
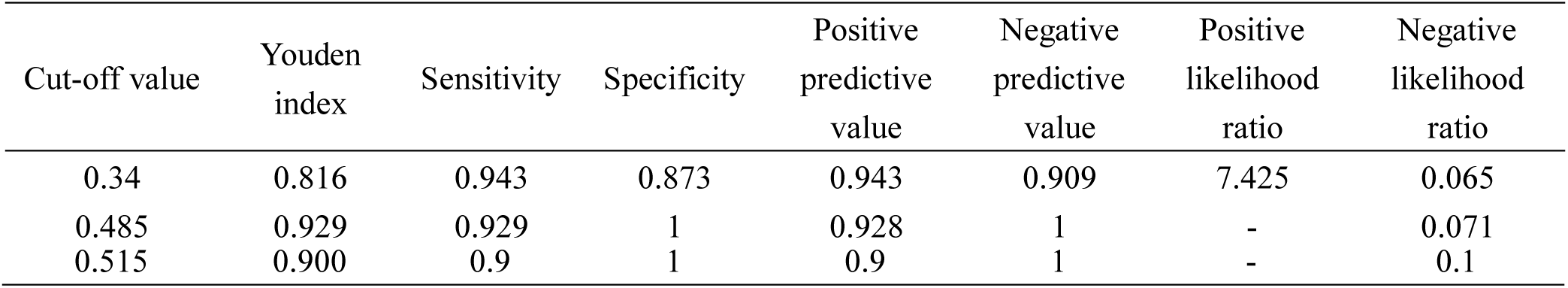
Diagnostic efficacy of the BALF GM test by cut-off value.

**Table 7.** Treatment status of patients in the case group.

| Project | Significant effect |  | Invalid | Total |
| --- | --- | --- | --- | --- |
|  | Cure | Better |  |  |
| Proven | 1 | 3 | 1 | 5 |
| Probable | 6 | 10 | 4 | 20 |
| Possible | 13 | 27 | 5 | 45 |
| Total | 20 | 40 | 10 | 70 |
| Proportion (%) | 86% |  | 14% | 100% |

## 3 Discussion

The incidence of pulmonary aspergillosis in patients with chronic respiratory diseases is increasing^[10]^. Because of its atypical clinical manifestations, pulmonary aspergillosis can be obscured by the symptoms of chronic respiratory diseases. Without treatment, the mortality rate increases significantly^[11]^, and patients with acute exacerbations of chronic respiratory diseases suffer cough and shortness of breath. Such patients may also have concomitant aspergillosis, which can be detected by chest imaging and is characterized by multiple plaques; when it cooccurs with IPA, the symptoms are likely to be confusing. Therefore, it is important to improve the diagnostic accuracy for chronic respiratory diseases cooccurring with IPA^[12]^. Pathogen culture takes some time to perform and has a low positive rate, so is of limited utility for early diagnosis of pulmonary aspergillosis. Furthermore, the decision for histopathological examination must be based on the patient’s tolerance and economic resources, limiting its clinical utility. GM, in serum, BALF, or cerebrospinal fluid is a biomarker of *Aspergillus*. The diagnostic efficacy of GM testing is high, and GM is now considered a diagnostic marker for aspergillosis^[13]^. Sulahian *et al*.^[14]^ reported that the results of GM tests are available 5–8 days earlier than those of imaging modalities. BALF can also be analyzed for *Aspergillus*, and BALF originates from the site of infection^[15]^. The positive rate of fungal cultures obtained by bronchoscopy is 46–75% ^[16, 17]^, and varies depending on disease course, disease severity, and whether antifungals have been administered.

Only seven patients in this study were positive for GM in serum; *Aspergillus* fungi did not invade the blood vessels, suggesting that these patients may still have been in the early stage of the disease. The serum GM-positive rate was relatively low, suggesting that blood and BALF GM tests may be more suitable for the diagnosis of chronic respiratory diseases and pulmonary aspergillosis. Meersseman *et al*.^[18]^ reported that among 25 patients with confirmed COPD and pulmonary aspergillosis, only 12 were positive for serum GM, but all were positive for BALF GM. In a prospective, single-center study^[19]^, among 110 patients with non-granulocyte deficiency, GM in BALF was more sensitive as an early diagnostic marker of pulmonary aspergillosis than GM in serum; the sensitivity values were 88% and 42%, respectively, with comparable specificity. At present, there is no standard cut-off value for the BALF GM test^[20-23]^. In a meta-analysis of the diagnostic performance of GM, Junling *et al*.^[24]^ showed that, with cut-off values of 0.5, 0.8, and 1.0, the sensitivity was 0.75, 0.69, and 0.68, and the specificity was 0.89, 0.94, and 0.96, respectively.

BALF GM cut-off values are set for patients with pulmonary aspergillosis with or without a variety of underlying diseases, and may not be suitable for patients with chronic respiratory disease^[25]^. Therefore, a classification system for patients with chronic respiratory diseases and pulmonary aspergillosis infection is needed, and a BALF GM cut-off value that is in line with clinical practice should be identified. This study included patients with chronic respiratory diseases, and assessed risk factors for the presence of *Aspergillus* infection and pulmonary lesions on imaging, Bronchoscopic examination and alveolar lavage were completed, and BALF-GM test of alveolar lavage fluid was carried out according to relevant diagnostic criteria. Patients clinically diagnosed with pulmonary aspergillosis were classified as cases, and patients without this diagnosis were controls. A total of 180 cases were included in this study; 70 in the case group and 110 in the control group. Statistical analysis of the BALF GM values showed that the mean rank of the case group was 140.80, which was significantly higher than that of the control group (58.49). Also, the BALF GM value of the case group was higher than that of the control group. Mann-Whitney U and Wilcoxon W-tests yielded a Z-value of -10.335 (P = 0.000), showing that the general distribution of BALF GM was significantly different between the two groups. The AUC was 0.957, indicating that BALF GM is an effective diagnostic marker for *Aspergillus* pneumocystis. According to the curve coordinate system and the Youden index, a GM cut-off value of 0.485 had the highest diagnostic efficacy for pulmonary aspergillosis accompanied by chronic respiratory diseases (sensitivity, 92.9%; specificity, 100%).

Early antifungal treatment of pulmonary aspergillosis can improve the prognosis. Seventy patients in this study were treated with antifungals (such as voriconazole and fluconazole). After treatment with antifungals, 20 cases were cured and 40 were improved (efficacy rate, 86%). Ten patients experienced adverse effects (adverse event rate, 14%), one of whom died. According to the 2016 guidelines on *Aspergillus* infection diagnosis and treatment^[6]^ of IDSA, oral antifungals can delay disease progression and improve symptoms and quality of life. Voriconazole is recommended as a first-line treatment. Isoconazole, amphotericin B liposomes, and other lipid preparations can be used as the second-line treatment. Echinomycin is not recommended as the first-line treatment. Huiqing *et al*.^[26]^ found that early empiric antifungal therapy significantly improved the prognosis compared with antifungal therapy after diagnosis. The antifungal treatment cycle should be ≥ 6–12 weeks, and should be determined according to the patient’s drug tolerance, immune status, clinical manifestations, laboratory test results, and imaging findings.

The diagnostic utility of GM in BALF for pulmonary aspergillosis is an important area of respiratory disease research. However, most prior studies focused on pulmonary aspergillosis in isolation. The BALF GM level is affected by various factors; in this study of patients with chronic respiratory diseases, the optimum BALF GM cut-off value for pulmonary aspergillosis was 0.485, which was lower than that reported by other studies^[27]^. There are many risk factors for pulmonary *Aspergillus* infection, such as neutropenia, organ transplantation, blood tumors, granular and non-granular deficiencies^[28, 29]^, organ and non-organ transplantation^[30, 31]^, and hematological tumors^[32, 33]^. the best cut-off value of BALF-GM in pulmonary Aspergillus patients with different risk factors should be set According to the results of BALF-GM. further increasing the diagnostic efficacy of BALF GM for pulmonary aspergillosis cooccurring with other conditions. For diagnosis of chronic respiratory diseases cooccurring with pulmonary aspergillosis, in addition to the BALF-GM test, the diagnostic efficacy of pathogens in BALF, GM in serum, and imaging modalities warrants further investigation.

In summary, Our study demonstrates that The BALF GM test has diagnostic utility for chronic respiratory diseases cooccurring with pulmonary aspergillosis. But For pulmonary Aspergillus patients, it is perhaps that best cut-off value of BALF-GM may be different according to different risk factors. More clinical data are needed to explore BALF-GM values in patients with pulmonary aspergillus in different immune conditions.

## Declarations

### Ethics approval and consent to participate

The study was approved by the Ethics Committee of Peking university Shenzhen Hospital.

### Consent for publication

All presentations of the study have consent to publish.

### Availability of data and material

The materials described in the manuscript are available from the corresponding author on reasonable request.

### Authors’ contributions

Data acquisition, analysis and interpretation: PX, GBL, CZ, JMM, WSD, Writing the article or substantial involvement in its revision: GBL, PX.

## Acknowledgments

The authors are indebted to the nurses of the bronchoscopy Room for making contributions to this study.

